# Validation of Functional Polymorphisms Affecting Maize Plant Height by Unoccupied Aerial Systems (UAS) allows Novel Temporal Phenotypes

**DOI:** 10.1101/2020.09.30.320861

**Authors:** Alper Adak, Seth C. Murray, Clarissa Conrad, Yuanyuan Chen, Steven Anderson, Nithya Subramanian, Scott Wilde

**Affiliations:** Department of Soil and Crop Sciences, Texas A&M University, College Station, TX 77843-2474; National Key Laboratory of Crop Genetic Improvement, Huazhong Agricultural University, Wuhan, China; Department of Environmental Horticulture, Institute of Food and Agricultural Sciences, Mid-Florida Research and Education Center, University of Florida, Apopka, FL, 32703

**Keywords:** Unoccupied Aerial System, High Throughput Phenotyping, Temporal SNP Effects

## Abstract

Plant height (PHT) in maize (*Zea mays* L.) has been scrutinized genetically and phenotypically due to relationship with other agronomically valuable traits (e.g. yield). Heritable variation of PHT is determined by many discovered quantitative trait loci (QTLs); however, phenotypic effects of such loci often lack validation across environments and genetic backgrounds, especially in the hybrid state grown by farmers rather than the inbred state preferred by geneticists. A previous genome wide association study using a hybrid diversity panel identified two novel quantitative trait variants (QTVs) controlling both PHT and grain yield. Here, heterogeneous inbred families demonstrated that these two loci, characterized by two single nucleotide polymorphisms (SNPs), cause phenotypic variation in inbred lines, but that size of these effects were variable across four different genetic backgrounds, ranging from 1 to 10 cm. Weekly unoccupied aerial system flights demonstrated both SNPs had larger effects, varying from 10 to 25 cm, in early growth while SNPs effects decreased towards the end of the season. These results show that allelic effect sizes of economically valuable loci are both dynamic in temporal growth and dynamic across genetic backgrounds resulting in informative phenotypic variability overlooked following traditional phenotyping methods. Public genotyping data shows recent favorably selection in elite temperate germplasm with little change across tropical backgrounds. As these loci remain rare in tropical germplasm, with effects most visible early in growth, they are useful for breeding and selection to expand the genetic basis of maize.

## Introduction

Plant height in maize has been subjected to many phenomic and genomic investigations since it influences plant architecture and agricultural performance, relating to other agronomically and economically significant traits in maize (*Zea mays* L.) (**Anderson *et al*. 2019, Farfan *et al*. 2013, Farfan *et al*. 2015, Sibov *et al*. 2003, Lima *et al*. 2006, Sari-gorla *et al*. 1999, Peiffer *et al*. 2014**). A key component of success to the Green-revolution, was the manipulation of plant height in wheat (*Triticum spp*.) and rice (*Oryza Sativa*) through the introduction of dwarf loci, initially used as a breeding strategy to maintain grain yield lost through lodging (**Peng *et al*. 1999, Khush 2001**). However, an important post-script has been that taller plant height leads to better yields in a number of cereal crops including rice (**Zhang *et al*. 2017**), sweet sorghum (**Shukla *et al*. 2017**), wheat (**Navabi *et al*. 2006**), and in maize (**Farfan *et al*. 2013**); as long as lodging can be avoided. Specifically, Farfan et al. (2013) evaluated that manual measured terminal plant height was positively correlated (r=0.61) with grain yield in overall subtropical environments. They proposed that an optimal taller plant height is a desirable maize ideotype with respect to yield, especially under heat and drought stress, as long as lodging is not an issue.

The wealth of studies on maize plant height have demonstrated the complexity, dynamic pattern and polygenic inheritance of this trait; genetically following the infinitesimal model, a trait governed by a large number of loci but with minor effects (**Wallace *et al*. 2016, Wang *et al*. 2019, Peiffer *et al*. 2014**). Thus far at least 219 QTLs have been identified as controlling the plant height in maize (http://archive.gramene.org/qtl/). Very few of these have been confirmed as QTL in independent studies across different genetic backgrounds and environments.

In contrast, the large effect genes identified with maize plant height have been associated with novel mutant alleles in hormone pathway genes; alleles rare or absent in landrace and elite cultivars because they are deleterious to plant fitness in nature. For instance, the dwarfing gene *dwarf 8* and *dwarf 9* encode DELLA proteins, which repress GA-induced gene transcriptions in the absence of GA signaling (**Lawit *et al*. 2010**); the *Dwarf3* gene (D3) of maize has significant sequence similarity to the cytochrome P450, which encodes one of the early steps in Gibberellin biosynthesis (**Winkler and Helentjaris 1995**); *brachytic2* mutants, the polar movement of auxins were hindered which resulted in compact lower stalk internodes (**Multani *et al*. 2003**), and *nana plant1* effects brassinosteroid synthesis (**Hartwig *et al*. 2011**).

That quantitative genetic loci for of plant height diversity still segregating in maize have not been cloned, let alone manipulated has likely been due to (i) limitations in detection ability of height related QTLs in diverse structure of mapping populations (**Xu *et al*. 2017**), (ii) aberrant plasticity under different plant architectures and genetic backgrounds (**Pigliucci 2005, El-soda *et al*. 2014**), (iii) aberrant plasticity under different environments, and genetic-by-environmental interactions (**Gage *et al*. 2017, El-soda *et al*. 2014**) and (iv) antagonistic pleiotropy of major genes (**Peiffer *et al*. 2014**). This is likely compounded by the use of inbred lines in genetic mapping as opposed to test-crossed hybrids, which tend to behave more stably, but also have less genetic variation. Maize evolved as a heterogenous and heterozygous outcrossing species and inbred lines expose weakly deleterious alleles uncommonly exposed in nature.

A genome wide association study (GWAS) on test-crossed hybrids made between a diversity panel and a line from the Stiff Stalk heterotic group (Tx714) under variable management discovered three significant loci associated with terminal plant height and yield by Farfan et al. (2015). These loci explained up to 5.6 cm per variant (4.6% of total), two of which also ranged from 0.14 to 0.59 ton/ha effects on grain yield (4.9% of total). Past GWAS studies have shown false positives due to cryptic population structure, familial relatedness, allele variants with low frequency or various allelic variants, as well as spurious associations between phenotypic variations and unlinked markers. For this reason, loci must be validated using different populations, environments (**Larsson *et al*. 2013**) and growth stages.

Although hybrid maize populations were used hardly ever in GWAS study, the identified genes by using hybrid populations in GWAS might be the key factor since candidate genes can be discovered under both overdominance and dominance genetic variances of heterosis, which fit better to be heterosis-related candidate genes (**Wang *et al*. 2017, Farfan *et al*. 2015**). This study for the first time (i) validated the temporal genes effects, which were discovered using hybrid genetic background in GWAS, in advanced heterogeneous inbred families (HIFs) generated from different parental crosses and (ii) implemented UAS platform to detect temporal changing of these genes’ effects on plant heights of HIFs.

## Materials and Methods

### Development of heterogenous inbred family (HIF) populations

Four linkage populations, segregating for the two SNPs of interest, were developed from crosses: (1) LH82 x LAMA, (2) Ki3 x NC356, (3) NC356 x Ki3 and (4) Tx740 x NC356 (recurrent parent x donor parent for populations 1 to 4) respectively, backcrossed with the recurrent parent. Individuals with heterozygous SNP calls (X:Y; i.e. donor allele: recurrent parent allele) were advanced; these seeds were planted in the following years nursery to be backcrossed (generally to BC_4-5_) or later selfed (generally to BC_4-5_F_3-4_). In the final selection step to develop HIFs within families, both SNPs (SNP1:SNP2) were selected in each population, both opposite (XX:YY and/or YY:XX) and identical (XX:XX and/or YY:YY) (**Fig. 1**).

**Figure 1.**
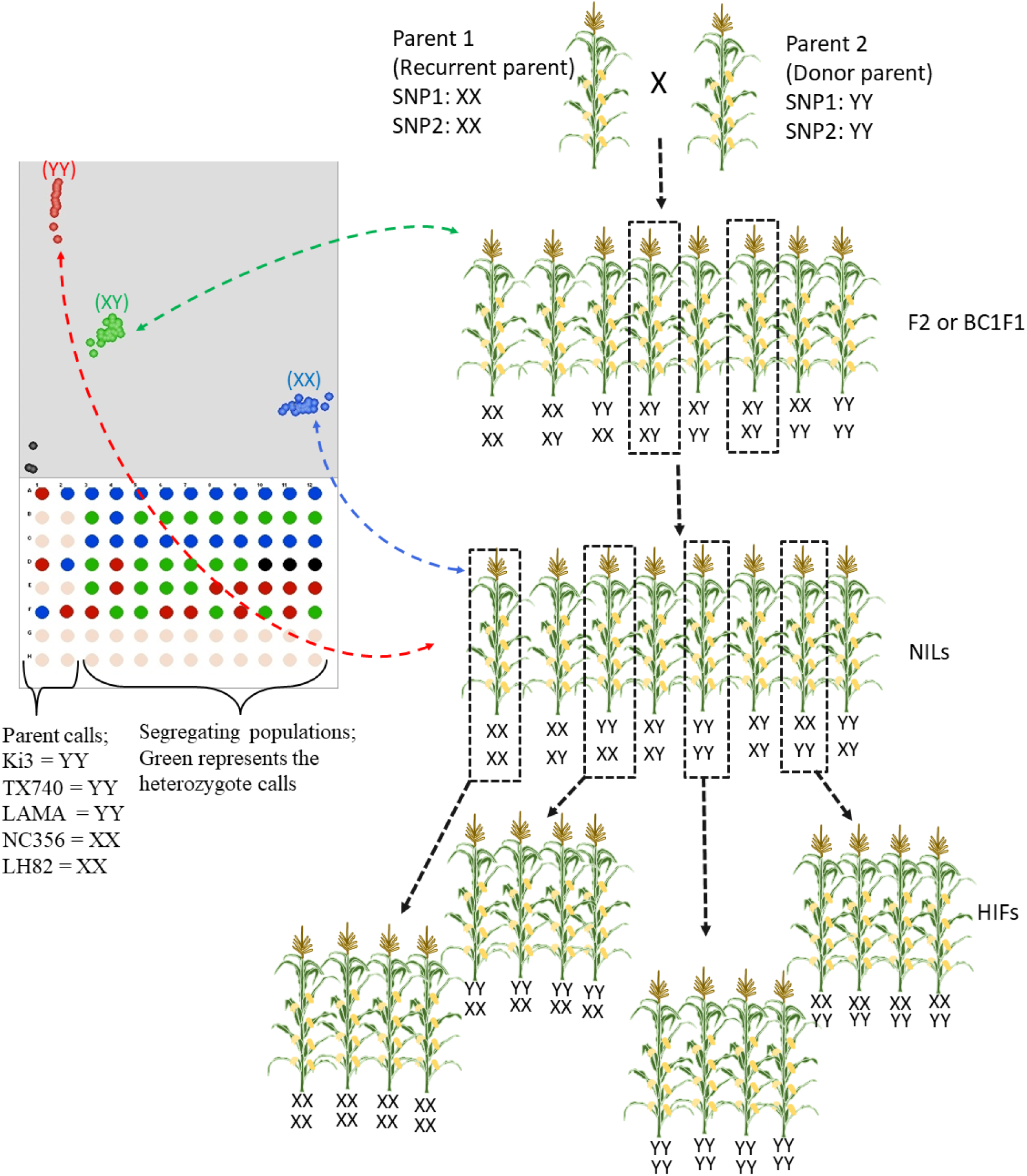
Breeding scheme of generating heterogeneous inbred families (HIFs) based on two SNP models and selection stages of pedigrees via KASP-PCR technology. † 10 to 20 plants from each plot were randomly selected. Of those having heterozygous loci (XY) were only selected and their ears were grown as rows (ear-to-row selection). After obtaining NILs, homozygous calls from both SNPs were selected as both identical (XX:XX, YY:YY) and opposite (XX:YY, YY:XX) to generate heterogeneous inbreds families. All parents were genotyped with pedigrees (left). Parents; Ki3, NC356, Tx740 and LH82, calls (SNP1:SNP2) are YY:YY, XX:XX, YY:YY and XX:XX respectively. No template controls, black color in figure, were used in each plate as negative controls.

### DNA extraction and KASP-PCR genotyping

Total genomic DNA was extracted from the frozen (−60**°**C) plant flag leaf tissue using a modified cetyltrimethylammonium bromide (CTAB) method (**Chen and Ronald 1999**). To design the unique markers targeting the SNP1 and SNP2, around 100 bp surrounding the two SNPs on either side were selected to determine allele-specific primers and allele general SNPs using BatchPrimer3 v1.0 (**You *et al*. 2008**). Sequence information of primers were obtained from (**Chen 2016**). Loci implemented into Kompetitive Allele Specific PCR (KASP) (http://www.kbioscience.co.uk/) assays by Chen (2016), were used in marker assisted backcrossing to develop HIFs across different near isogenic line (NIL) backgrounds. and were used to detect SNP calls (XX, XY and YY) for developing HIFs during 2016 to 2019 (**Fig. 1)**.

Tassel software (version 5) (**Bradbury *et al*. 2007**) was used to obtained linkage disequilibrium (LD) (LD windows size = 50 markers) to find the nearby LD patterns of SNPs locations. MaizeGBD (http://www.maizegdb.org/) genome browser was used to determine the genes linked to SNPs. The Gramene database (http://www.gramene.org) was used for the identification of candidate genes. The Panzea (https://www.panzea.org/) website was used to extract the sequence information of genes from publicly available maize germplasm. The years information when germplasm was developed were obtained from Germplasm Resources Information Network (GRIN, https://www.ars-grin.gov).

### Planting and agronomic practices

Plants were grown near College Station, TX (coordinate: 30°33’00.8” N 96°26’04.3” W) for summer nurseries and Weslaco (26°09’32.7” N 97°57’36.1” W), Texas for winter nurseries from 2016 to 2019. Most of these nurseries used small sample sizes of as little as one plot for per plot (including X:Y) to primarily advance and increase. For phenotyping in College Station 2019, entire plots of X:X, Y:Y, and X:Y for each HIF were planted on the 12^th^ of April, 2019, in two replicates; although there were often more than one representative for each genotype. In total, 288 plots were grown. These were planted in a split:split:split plot design where the main split was replicate, the second split was population / genetic background, and the third split was genotype. Unless noted, all reported hand measurements and UAV flights were conducted when HIFs were grown near College Station in 2019.

### Phenotyping

Days to anthesis (DTA) and silking (DTS) were recorded on a plot basis when 50% of the plants were showing anthers and silks respectively, checking plots daily. Three different terminal plant height measurements were taken using a ruler including TH, FH, EH July 2^nd^, 2019, about two to three weeks after flowering. In addition, unoccupied aerial vehicle (UAV, aka drone) plant height measurements were taken weekly from emergence to the end of the growth period. The flight dates were shown as day/month/year (dd/mm/yy).

UAV images of the field were taken using a DJI Phantom 4 Pro V2.0 (DJI, Shenzhen, China) at an above ground altitude of 25 meters. The standard integrated camera resulted in images having a resolution of 72 DPI. DJI standard flight control software was used. Orthomosaics and point clouds were created with the images for each flight by using Agisoft Metashape V15.2 software (Agisoft LLC, Russia). The captured images were at 72dpi with 90 percent overlap and were used to create an orthomosaic and point cloud for determining the plant height as a function of time during the growth period. Ground control points (GCPs) were used during the flights to assist the data processing and reduce effects due to aberrations and the resulting georeferenced mosaics.

Previous work has shown that various methods to measure inbred maize plants from the ground using point clouds produced similar results (**Anderson *et al*. 2020**). Point clouds of each flight were processed using CloudCompare (version: 2.11. alpha). To set a canopy height model (CHM), first flight containing bare ground was used as a digital terrain model (DTM). Digital surface model (DSM) of each flight was subtracted from DTM to calculate CHM (**Fig. S1)**. Each plot was drawn using the polygon function of CloudCompare.

### Statistical inference

Statistical models were developed according to the distribution of SNP1 and SNP2 combinations obtained from the HIFs. Spatial variation was partitioned as random effects into ranges and rows. Each model was run using a restricted maximum likelihood (REML) method in JMP version 15.0.0 (SAS Institute Inc., Cary, NC, USA) to predict the best linear unbiased estimates (BLUEs) of SNPs. In these models, fit as fixed effects to obtain BLUEs values during flights as well as for ruler measurements; as well as fit as random effects in an all random model to obtain variance components. All components, except the SNPs, were always fit as random effects under the following mixed linear models (MLM) in each model.

First, each SNP was tested separately within each population (Equation [1]). While one of two SNPs was segregating, the other one was fixed (not segregating as XX or YY) in respective populations to compare the BLUEs of SNP calls. This equation was used for hand measurement data on a plant basis for each population.

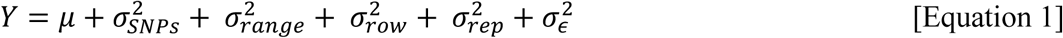

Within this base model, response variable (*Y*) was one of the three hand measures of plant height data; 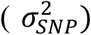 represented variance of one of SNPs to be tested on condition that other one is fixed XX and/or YY within each respective population. Other variance components, including range 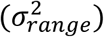, row 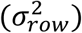 and rep 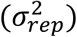, account for the spatial variation. 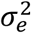 is the pooled unexplained residual error.

Plant height and flowering time were also tested for SNP1 and SNP2 individually combining all data across populations 1, 2 and 3 (Equation [2]). While one of the two SNPs segregated, the other one was fixed (not segregating as XX) in the model. In this equation, the population 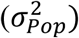 effect was added compared to Equation [1]. BLUEs and BLUPs of SNPs and their interactions with populations respectively were obtained for each UAS flight and ruler measurement.

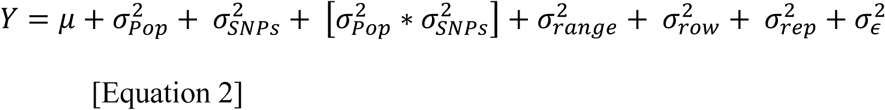

The interactions of both SNPs and populations using the full factorial function was tested for both flowering time and for plant height from the ruler measurement and UAS flights temporally across populations 1 and 2 (Equation [3]).

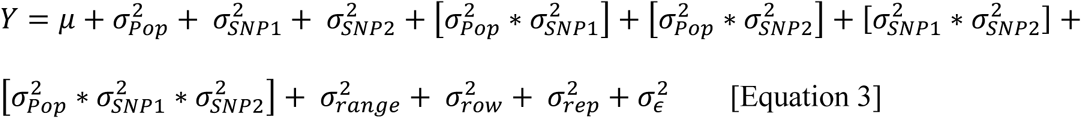

In here, response variable (*Y*) is plant height data. 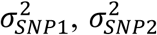 and 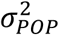 represent the variance components of SNP1, SNP2 and population respectively while other variance components were the same as stated previously in Equation [1] and Equation [2]. In this equation population 1 and 2 were used.

Repeatability (R) was calculated based on following formula with number of replication (r) for single environments (Equation [4]).

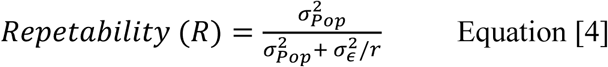

Additional data processing and visualizations were performed in R version 3.5.1 (**R core team 2018**).

## Results

The effects of Cytosine/C for SNP1, Adenine/A for SNP2 (e.g. XX) calls in both SNPs, contributed by both NC356 and LH82 parents **(Fig. S2)**, increased all three measures of plant heights (TH; from ground to tip of tassel, FH; from the ground to the flag leaf collar, EH; first ear height from the ground to first ear shank). Tassel height differences between XX and YY calls were statistically significant across all populations (**Fig. 2**), varying from 2.0 to 8.9 cm (SNP1) and 3.0 to 11.9 cm (SNP2) depending on the populations genetic background (**Fig. 2**).

**Figure 2.**
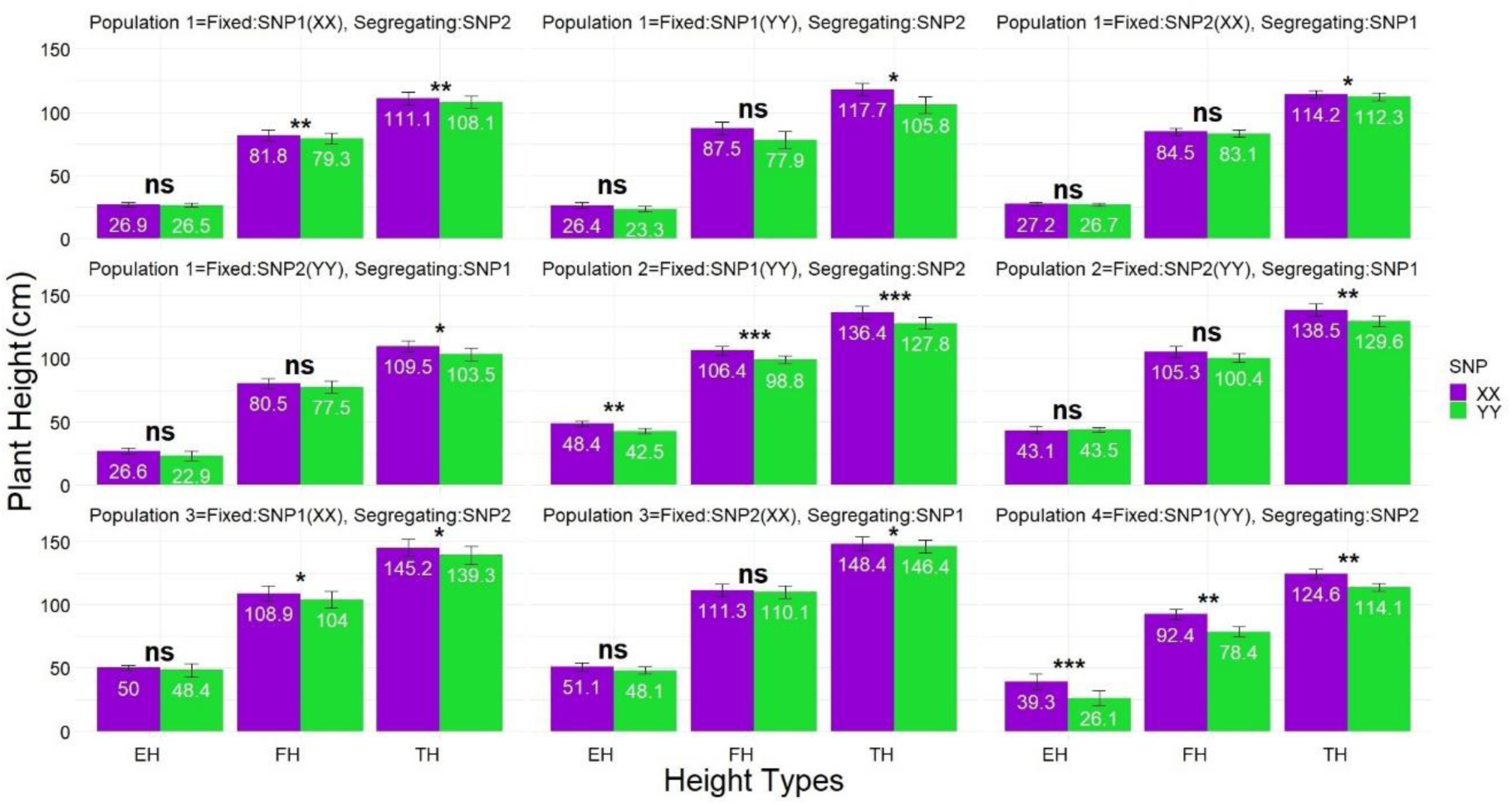
BLUEs showed XX calls significantly increased all three ruler measures of plant heights in a consistent direction across populations. † Population 1, 2, 3 and 4 represents near isogenic lines of [LAMA (recurrent parent) x LH82], [Ki3 x NC356 (recurrent parent)], [Ki3 (recurrent parent) x NC356] and [Tx740 (recurrent parents) x NC356] respectively. ‡ Best linear unbiased estimators (BLUEs) were calculated using equation 1 (Equation [1]). *, **, *** indicate significance level at 0.05, 0.01 and 0.001 respectively while ns indicate not significant. § TH, tip of tassel height; FH, flag leaf collar height; and EH, height of the first ear shank from ground on the x-axis

The favorable locus (XX) of SNP1 and SNP2 across populations increased TH ∼ 4 cm and FH ∼ 3 cm (Equation [2]; **Fig. S3**). Interactions between SNP1*population and SNP2*population varied with TH differences were observed up to 10 cm, followed by up to 7.0 cm for FH (**Fig. S4)**.

Flowering times (days to anthesis, DTA, and days to silk, DTS) when used as response in Equation [2] demonstrated the taller XX allele of SNP1 and SNP2 for plant heights also caused later flowering. XX allele of SNPs delayed flowering times between 1 and 5 days depending on the genetic backgrounds of populations (**Fig. S5**). Result of orthogonal contrasts conducted between calls of each population showed this lateness was statistically significant (**Fig. S5**).

In Equation [3], SNP1 and SNP2 interaction 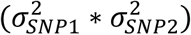 for TH and combined interaction with populations 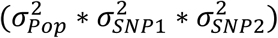 were found to be significantly taller than shortest combination (YY-YY) when either SNP1, SNP2 or both were XX favorable locus, resulting in that combined favorable SNP1 and SNP2 loci (XX-XX) was tallest in TH, which was 8.8 cm taller than YY-YY combination (**Fig. S6**). This was 3.5 cm taller than expected from SNP1 or SNP2 alone and represents a synergistic effect between these two loci. There was also an epistatic effect of these SNPs with XX-XX combination increasing height 8 cm in population 1 but 9.6 cm for population 2 and was consistent for other measurements of plant height (**Fig. S7**).

The proportion of total experimental variance attributable to differences between populations 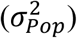 varied from 64 to 80 percent within Equation [2] and Equation [3] for plant height measurements by ruler. Population effects, spatial (range, row) partitioned large amounts of experimental variance, but repeatability was high at 89 to 95 percent (**Table S1 and S2**).

### Statistical inferences of UAS PHT

Temporal resolution of each UAS flight captured that the highest plant height (Crop Height Model; CHM) differences between favorable (XX) and unfavorable loci (YY) were 16 to 20 cm in early growing stages (34 to 54 days after sowing; first four flights) but narrowed 3 to 5 cm by harvest time depending on when either SNP1 or SNP2 were tested in Equation [2] respectively (**Fig. 3**). The differences between favorable and unfavorable loci varied depending on interaction between population with SNP1 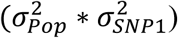 and population with SNP2 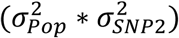 by Equation [2]. The differences between calls in either interactions had a descending pattern from early growing season to time of harvest, showed that highest differences between calls for populations were captured between 9 to 26 cm in early season and narrowed 1 to 10 cm by the time of harvest (**Fig. 4**).

**Figure 3.**
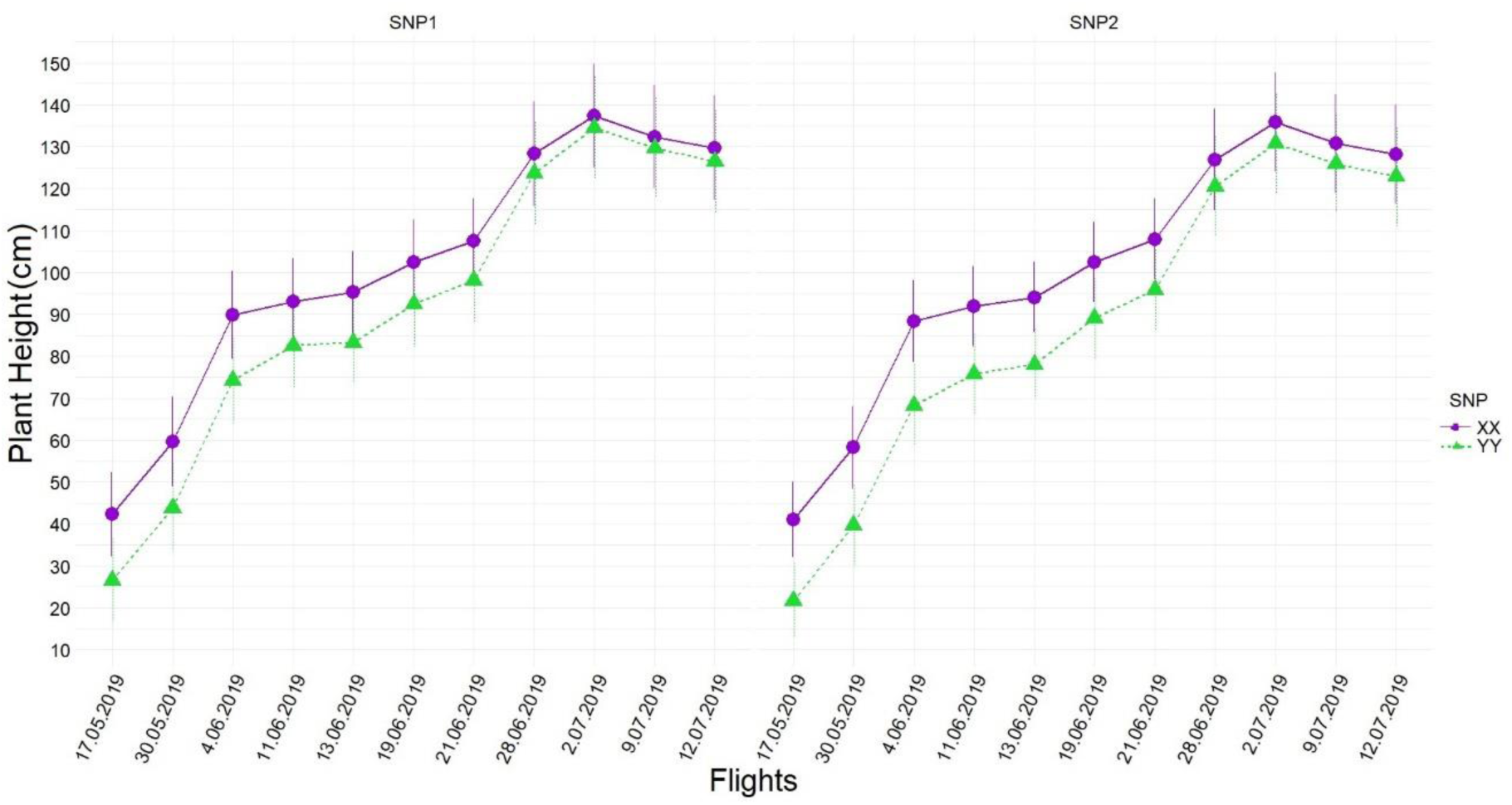
Temporal resolution of differences between SNP1 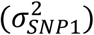 (left) and SNP2 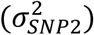 (right) calls obtained by Equation [2] during UAS flights across all populations.

**Figure 4.**
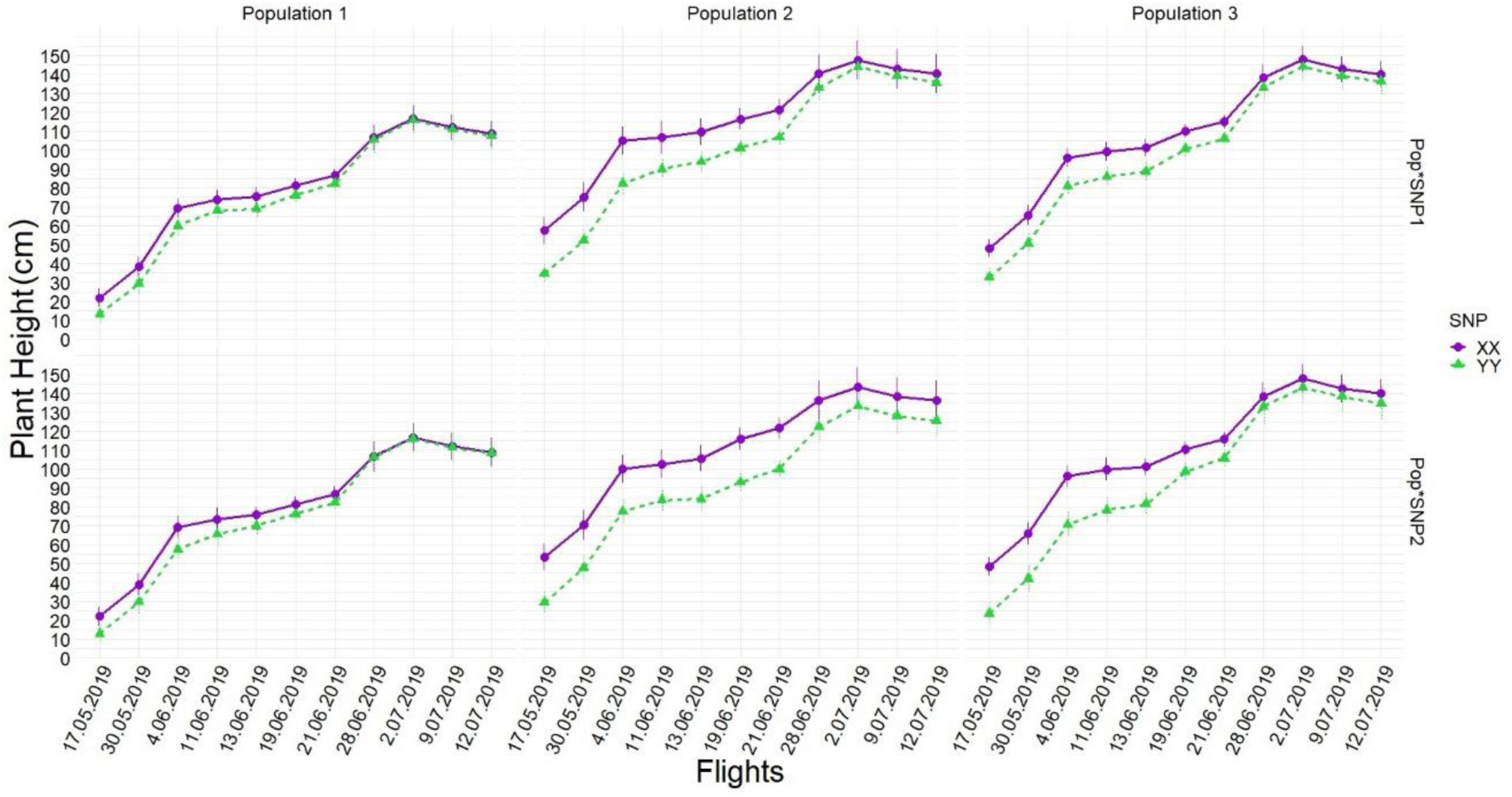
Temporal resolution of interactions of 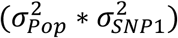 and 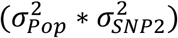 obtained by Equation [2] during UAS flights showed large differences between how the SNPs behaved on different genetic backgrounds.

In Equation [3], UAS captured that favorable loci combination of XX-XX (SNP1:SNP2) was found tallest in every flight followed by YY-XX, XX-YY and YY-YY (**Fig. 5**), resulting in that height differences between favorable and unfavorable loci combination for population 1 and population 2 of 11 to 25 cm in the early growing stages and 7 to10 cm by the time of harvest (**Fig. 6**). Synergetic effect of favorable loci combination on unfavorable loci combination also decreased from 9 cm to 2 cm as the growing period progressed.

**Figure 5.**
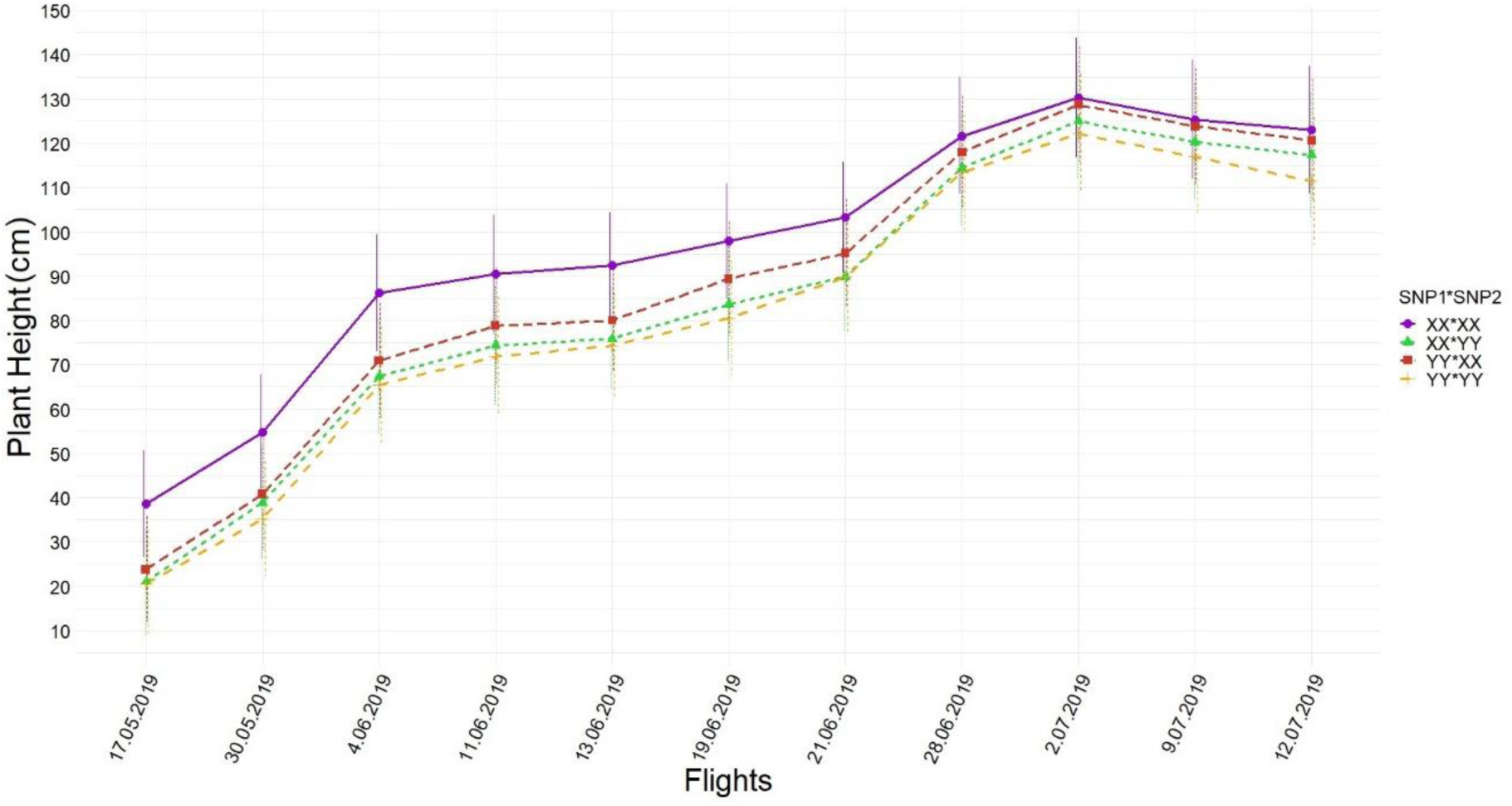
Temporal resolution of differences among SNP1-SNP2 interactions 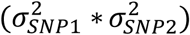 obtained by Equation [3] during UAS flights show a synergistic effect on increasing height.

**Figure 6.**
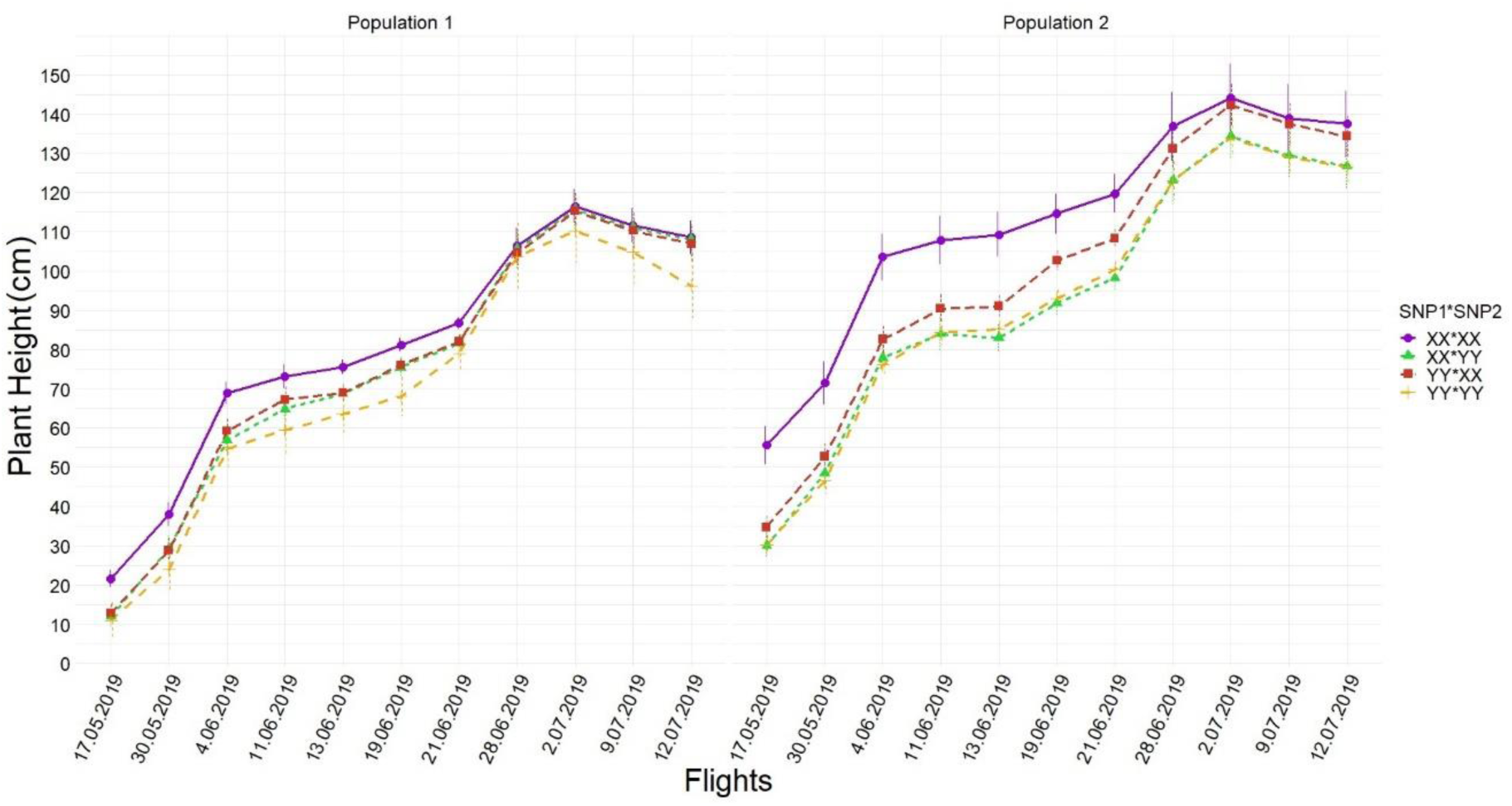
The SNP combinations work differently across different populations genetic backgrounds and over time. Using 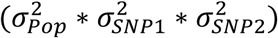 obtained by Equation [3] during UAS flights

Population variation 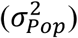 always explained the highest percentage of total variation in both Equation [2] and Equation [3], resulting in repeatability estimates which fluctuated between 84 and 97 percent (**Table 1 and 2**) during growing periods for plant height. SNP1 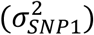 and SNP2 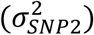 in Equation [2] showed decreasing trends from ∼20-30 percent of explained total variation to below 1 percent during growing periods (**Table 1**) as well as decrease from ∼2-5 percent to below 1 percent in the interaction of SNPs in Equation [3] (**Table 2**)

**Table 1.** Percentages of total variance explained by each compoenets in Equation [2] when SNP1 was tested (above) and SNP2 was tested (below) as well as the total variance in number and repeatibilty for each UAS flight.

| Variance component<br>(Random effect) | Percentage of variation explained by each variable component for each flight |  |  |  |  |  |  |  |  |  |  |
| --- | --- | --- | --- | --- | --- | --- | --- | --- | --- | --- | --- |
|  | <u>17.05.19</u> | <u>30.05.19</u> | <u>4.06.19</u> | <u>11.06.19</u> | <u>13.06.19</u> | <u>19.06.19</u> | <u>21.06.19</u> | <u>28.06.19</u> | <u>2.07.19</u> | <u>9.07.19</u> | <u>12.07.19</u> |
| Population | 45.7 | 46.2 | 45.5 | 47.1 | 47.2 | 64.3 | 66.0 | 54.0 | 53.8 | 54.3 | 54.1 |
| SNP1 | 20.4 | 18.1 | 18.9 | 9.1 | 13.9 | 8.1 | 7.6 | 0.7 | 0.3 | 0.3 | 0.4 |
| Population*SNP1 | 2.6 | 1.8 | 1.7 | 1.7 | 1.8 | 1.1 | 1.3 | 0.0 | 0.0 | 0.0 | 0.0 |
| Replication | 8.0 | 9.4 | 8.5 | 7.4 | 7.8 | 4.9 | 4.7 | 14.7 | 14.5 | 14.2 | 14.0 |
| Row | 0.2 | 0.3 | 0.0 | 1.1 | 0.5 | 0.5 | 0.6 | 3.4 | 3.2 | 3.3 | 3.2 |
| Range | 11.7** | 13.0** | 13.8** | 14.3** | 15.9** | 12.4*** | 10.9** | 7.0* | 7.5* | 7.2* | 7.7* |
| Residual | 11.4 | 10.8 | 11.7 | 19.3 | 12.9 | 8.7 | 9.0 | 20.3 | 20.7 | 20.7 | 20.6 |
| **Total variation in number** | 449.4 | 490.1 | 476.7 | 474.9 | 412.5 | 395.4 | 371.8 | 547.8 | 551.3 | 550.1 | 559.3 |
| **Repeatability (R)** | 0.89 | 0.89 | 0.87 | 0.83 | 0.88 | 0.94 | 0.94 | 0.84 | 0.84 | 0.84 | 0.84 |
| Variance component<br>(Random effect) | Percentage of variation explained by each variable component for each flight |  |  |  |  |  |  |  |  |  |  |
|  | <u>17.05.19</u> | <u>30.05.19</u> | <u>4.06.19</u> | <u>11.06.19</u> | <u>13.06.19</u> | <u>19.06.19</u> | <u>21.06.19</u> | <u>28.06.19</u> | <u>2.07.19</u> | <u>9.07.19</u> | <u>12.07.19</u> |
| Population | 30.9 | 32.3 | 32.8 | 34.7 | 32.2 | 50.9 | 88.2*** | 48.4*** | 50.5*** | 49.2*** | 82.0*** |
| SNP2 | 32.4 | 27.6 | 30.8 | 21.9 | 24.2 | 16.6 | 0.1*** | 0.1*** | 0.2*** | 0.1*** | 0.1*** |
| Population*SNP2 | 7.1 | 5.8 | 3.9 | 4.3 | 7.9 | 7.1 | 0.0 | 0.1*** | 0.1*** | 0.1*** | 0.1*** |
| Replication | 9.2 | 11.3 | 9.3 | 12.2 | 7.8 | 6.2 | 0.4 | 30.4 | 27.3 | 28.0 | 7.2 |
| Row | 0.1 | 0.1 | 0.1 | 0.1 | 0.2 | 0.8 | 0.7 | 0.8*** | 0.6*** | 0.4*** | 1.0*** |
| Range | 11.9** | 14.3** | 14.8** | 17.2** | 16.3** | 11.8** | 5.7*** | 7.4*** | 7.3*** | 7.3*** | 2.7*** |
| Residual | 8.4 | 8.6 | 8.2 | 9.7 | 11.3 | 6.6 | 4.9 | 12.9 | 14.0 | 14.9 | 6.8 |
| **Total variation in number** | 475.2 | 512.6 | 548.9 | 473.8 | 394.7 | 403.0 | 385.2 | 660.4 | 484.1 | 608.2 | 1379.2 |
| **Repeatability (R)** | 0.88 | 0.88 | 0.89 | 0.88 | 0.85 | 0.94 | 0.97 | 0.88 | 0.88 | 0.87 | 0.96 |

**Table 2.** Percentages of variances explained by each compoenets in Equation [3] as well as total variance and repetability for each UAS flights.

| Variance component<br>(Random effect) | Percentage of variation explained by each variable component for each flight |  |  |  |  |  |  |  |  |  |  |
| --- | --- | --- | --- | --- | --- | --- | --- | --- | --- | --- | --- |
|  | <u>17.05.19</u> | <u>30.05.19</u> | <u>4.06.19</u> | <u>11.06.19</u> | <u>13.06.19</u> | <u>19.06.19</u> | <u>21.06.19</u> | <u>28.06.19</u> | <u>2.07.19</u> | <u>9.07.19</u> | <u>12.07.19</u> |
| Population | 81.4*** | 81.1*** | 74.6*** | 81.1*** | 79.3*** | 68.3 | 84.8*** | 70.8*** | 57.4*** | 57.3*** | 57.4*** |
| SNP1 | 2.2*** | 2.7*** | 2.1*** | 1.5*** | 1.6*** | 1.4 | 0.1*** | 0.6*** | 0.7*** | 0.6*** | 0.3*** |
| Population*SNP1 | 0.1*** | 0.1*** | 0.1*** | 0.1*** | 0.1*** | 0.1 | 0.1*** | 0.3*** | 0.5*** | 0.6*** | 0.6*** |
| SNP2 | 5.5*** | 3.9 | 5.7 | 3.4 | 3.2 | 4.1 | 0.3*** | 2.0*** | 1.2*** | 1.0*** | 0.2*** |
| Population*SNP2 | 0.1*** | 0.2*** | 0.2*** | 0.1*** | 1.6*** | 6.1 | 3.5*** | 1.4*** | 2.2*** | 2.6*** | 2.2*** |
| SNP1*SNP2 | 0.7*** | 0.7*** | 1.2*** | 1.1*** | 1.8*** | 2.0 | 1.7*** | 0.2*** | 0.3*** | 0.3*** | 0.7*** |
| Population*SNP1*SNP2 | 0.7*** | 0.2*** | 0.1 | 0.1*** | 1.1*** | 0.1*** | 0.1*** | 0.1*** | 0.1*** | 0.1*** | 0.1*** |
| Replication | 0.7 | 1.4 | 1.3 | 1.3 | 0.0 | 0.0 | 0.0 | 7.4 | 5.1 | 4.8*** | 5.2 |
| Row | 0.2*** | 0.1*** | 0.1 | 0.1*** | 0.5*** | 1.1 | 0.9*** | 2.3*** | 2.9*** | 2.8*** | 2.8*** |
| Range | 3.3*** | 5.0*** | 7.2** | 5.2*** | 3.7*** | 6.4* | 2.4*** | 4.2*** | 11.2*** | 11.2*** | 11.5*** |
| Residual | 5.1 | 4.5 | 7.4 | 6.0 | 7.1 | 10.3 | 6.1 | 10.7 | 18.3 | 18.7*** | 19.1 |
| **Total variation in number** | 640.1 | 807.7 | 609.1 | 895.6 | 691.2 | 377.0 | 593.0 | 871.2 | 473.8 | 463.0 | 466.0 |
| **Repeatability (R)** | 0.97 | 0.97 | 0.95 | 0.96 | 0.95 | 0.93 | 0.96 | 0.93 | 0.86 | 0.86 | 0.86 |

### Accuracy assessment between UAS-PHT and TH

For accuracy assessment, means and medians of each plot measured by ruler on July 2^nd^, 2019 were correlated with UAS-PHT captured on the same date, and a correlation coefficient was found to be 0.83 for either the median or mean correlated with UAS-PHT **(Fig. S8)**.

### Genes colocalized with associated SNPs

Physical locations of both SNPs (SNP1 and SNP2 overlap GRMZM2G035688 and GRMZM2G009320) from Farfan et al. (2015) updated to genome version 5 showed that regions of LD blocks spanned around 48 bp (*D*^′^:1, sig = 0.00) for SNP1 and 878 bp (*D*^′^:1, sig =0.01) for SNP2 (**Fig. S2**).

## Discussion

These results demonstrated in maize for the first time that quantitative height loci first discovered through GWAS testcrossed diversity panel studies also conferred effects across four very diverse genetic backgrounds. An uncommonly discussed advantage of GWAS over linkage mapping is the ability to detect alleles that function non-specifically across genetic backgrounds; that is maximizing discovery context-independent alleles unaffected by genetic background epistasis that has hindered use of quantitative loci in the past. These alleles were first confirmed in linkage mapping populations (F_3:4_) developed from parental lines segregating for the two SNPs of interest (**Chen 2016**). However, Chen (2016) estimated different absolute effect sizes for these SNPs compared to those estimated in the initial GWAS study (**Farfan *et al*. 2015**).

Across thousands of studies, many maize loci have been associated with agronomic traits in maize (**Andersen *et al*. 2005, Farfan *et al*. 2015, Larsson *et al*. 2013, Li *et al*. 2013, Peiffer *et al*. 2014, Thornsberry *et al*. 2001, Weng *et al*. 2011, Anderson *et al*. 2018**). Although strong population structure and relatedness has been controlled in most GWAS studies to reduce false positive results (**Lipka *et al*. 2015, Myles *et al*. 2009**), we are cautioned by the cryptic population structure of *dwarf8* (**Larsson *et al*. 2013**) and possibilities of overfitting GWAS models to identify non-causal loci. Confirmation of loci from GWAS studies is therefore necessary to understand if the alleles are robust and useful as well as if the and effect sizes are consistent. Therefore, it is critical that the two SNPs used in this study were validated over HIFs from four linkage populations, as contributing to taller plant heights in both ruler measurements and UASs data.

### Temporal resolutions of SNPs effects on PHT

The first seven UAS flights, flown during vegetative growth (typically up to 70 days after planting), found the largest SNP effect sizes and SNPs interaction effects (**Fig. 3, 4, 5 and 6**) as well as explained the most variation **(Table 1 and 2**). This was surprising since these SNPs were initially discovered in the GWAS panel through terminal height measurements using a ruler (**Farfan *et al*. 2015**); although UAS phenotyping technologies were not available when that study was conducted, and temporal ruler measurements would have been infeasible. The last four UAS flights were flown in the reproductive stage (days 70 to 100 after sowing) after vegetative growth period of increasing internodes had ended when the effect size of both SNP’s and their interactions had become much smaller, in agreement with ruler measurement results taken July 2^nd^, 2019 (82th day after sowing, between R5 to R6) **(Tables 1 and 2; Tables S1 and S2)**. However, in the reproductive growth phase, measuring plants individually with a ruler and plots by UAS, the differences between the main SNP effects can still be resolved **(Table S1 and S2)**. Maize yield has been most strongly correlated with plant height, in V6 (6-leaf), V10 (10-leaf) and V12 (12-leaf) growth stages, with V10 and V12 growing stages more important than other stages when earliness was desired (**Yin *et al*. 2011**). While little knowledge exists at intermediate growth time points, the evidence is much stronger between terminal plant height and grain yields (**Anderson *et al*. 2019**). Context-dependency effects of SNPs under different genetic backgrounds were also able to be resolved best in early UAS flights with larger effects sizes for populations 1 and 2 in the earliest flights (**Fig. 4 and 6**). Population 3, developed as a reciprocal cross of population 2, was also observed to have effect size differences (**Fig. 4**).

Recent studies showed the temporal plant height-SNPs associations in sorghum (**Miao *et al*. 2020**) and maize (**Anderson *et al*. 2020)**, however there were no precise validation of gene effects in either early or late growth stages. In conclusion, in this study, high-throughput phenotyping technology firstly enabled the monitoring temporal shift of gene effects with high resolution during growth periods, which cannot be captured to such an extent by traditional terminal measurement methods.

### Pleiotropy of SNPs with flowering times

Both SNPs had pleiotropic effects on flowering (**Fig. S5**) not observed in the initial GWAS study (**Farfan *et al*. 2015**). This was likely because heterosis in hybrid backgrounds tends to reduce or compress variation seen in inbred lines and because heterosis causes maize to flower earlier. Here the earliest flowering population had the least difference between alleles (pop 1, <0.5 days) while the latest flowering population was able to discriminate the largest differences (pop3, >2 days) (**Fig. S5**).

### Functional annotation of candidate genes

SNP1 (GRMZM2G035688) corresponded *aberrant phyllotaxy1* (also known as abph1) was first observed in maize mutant showing transformed phyllotaxy behavior (**Jackson and Hake 1999**). Phyllotaxy is the geometric arrangement of leaves and flowers to control the plant formation by shoot apical meristem (SAM). Unlike auxin action in phyllotaxy regulation in Arabidopsis (*Arabidopsis thaliana*), cytokinin-inducible type A response regulator is encoded by *abph1*, indicating that cytokinins play a role on aberrant phyllotaxy in maize (**Lee *et al*. 2009**). Auxin or its polar transport is necessity for *abph1* expression due to fact that *abph1* expression was dramatically lessened after treatment of a polar auxin transport inhibitor to maize shoots (**Lee *et al*. 2009**). Taken together, GRMZM2G035688 encoding *abph1* is essential for adequate maize PINFORMED (*PIN1)* expression, which is polar auxin transporter for leaf primordia expression in maize, and auxin localization in embryonic leaf primordia in SAM (**Lee *et al*. 2009**). SNP2 (GRMZM2G009320) encodes a Glyceraldehyde-3-phosphate dehydrogenase (GAPDH) which catalyze sixth step of glycolysis into energy as well as carbons in higher plants. Under stress conditions such as salt or oxidative stresses, the activity of enzyme increases to manipulate energy formation in plants (**Bustos *et al*. 2008, Zhang *et al*. 2011**).

### Recent breeding has selected the favorable allele

Previously, several genes important in post domestication adaptation were identified by comparing the maize lines from different early and late eras to show the proof of directional selection (**van Heerwaarden *et al*. 2011**) where our genes (GRMZM2G035688 and GRMZM2G009320) were not included. Recent publicly available genotyping of diverse public inbred lines and germplasm (∼1000 subset) for SNP1 and SNP2 allele information was extracted and grouped into five categories (**Fig. 7**) and compared by year of development or release. The frequency of SNP favorable alleles (X:X; increased height yield and flowering) showed consistent increases in most groups (**Fig. 7**). Expired plant variety protection lines (Ex-PVP) developed and released by industry and U.S. public lines showed the greatest shifts towards the favorable alleles, almost to fixation. A lower frequency but less dramatic shift in CIMMYT originated tropical lines suggests that these loci are still segregating in tropical maize germplasm perhaps because the effects are less dramatic in the tropics. So, these alleles are examples of favorable allele selection during time, especially in temperate areas, unsurprising given their large phenotypic effects.

**Figure 7.**
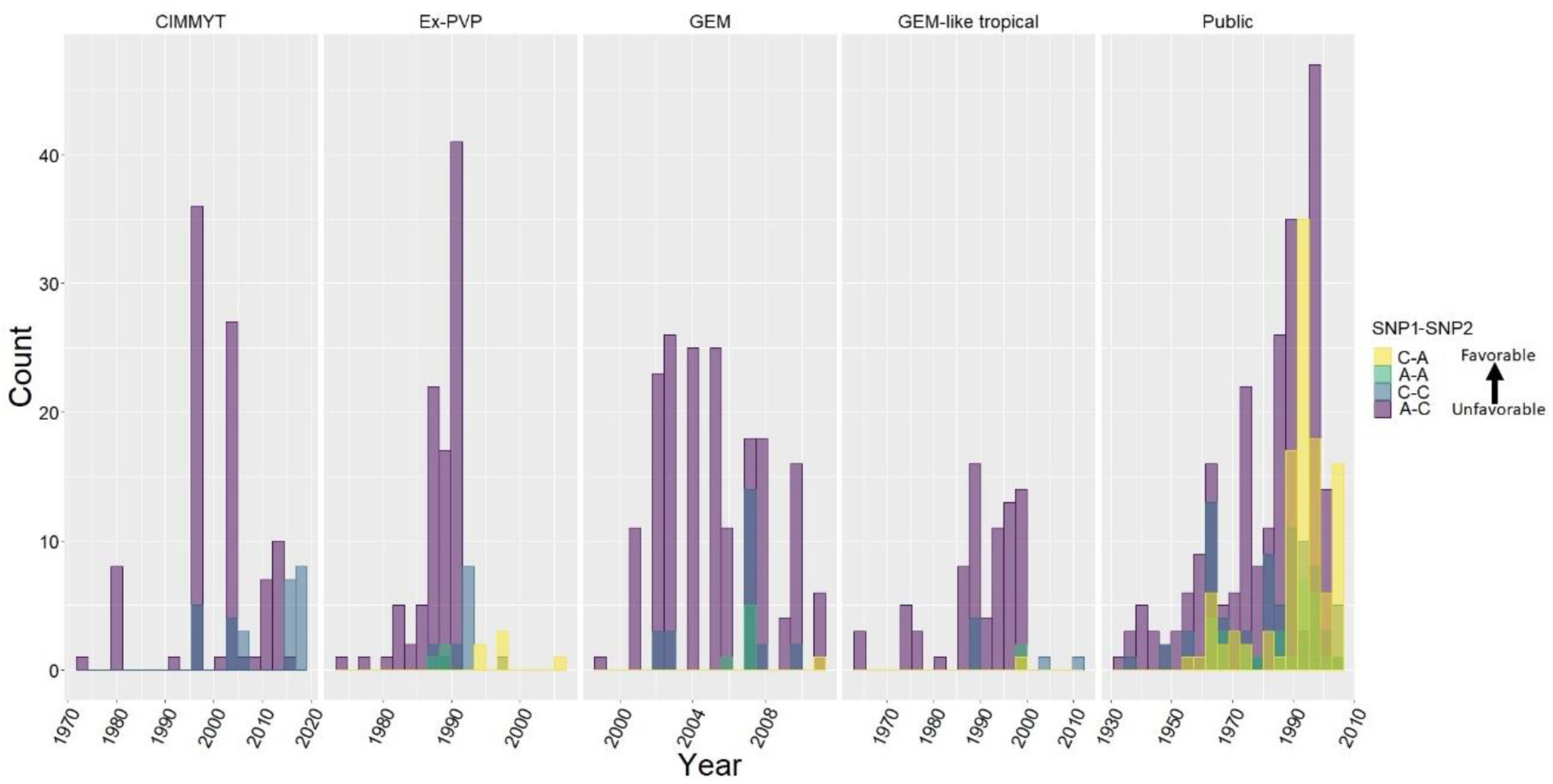
The allele combinations of SNP1 and SNP2 were illustrated in years and five germplasm categories.

In summary, a GWAS field study of hybrids under stress successfully nominated QTVs that work across genetic backgrounds, in inbred lines and throughout diverse environments. New UAS tools provided substantially more information and better screening for their effects than the traditional terminal ruler height measurements in which they were discovered. To get a better understanding of QTV’s affecting complex traits such as plant height and grain yield in maize, combination of high-throughput phenotyping and genotyping studies must be evaluated together, which will be critical for managing the phenotypic plasticity of complex traits.

## Acknowledgments

All authors declare that there is no conflict of interest. This study was conducted through financial support of USDA–NIFA–AFRI Awards 2017-67013-26185 and 2010-85117-20539, USDA–NIFA Hatch funds, Eugene Butler Endowed Chair, Texas A&M AgriLife Research, the Texas Corn Producers Board. Alper Adak was supported by a fellowship from Republic of Turkey, Ministry of National Education and Ministry of Agriculture and Forestry.

## Supplementary Figures and Tables

**Fig. S1.**
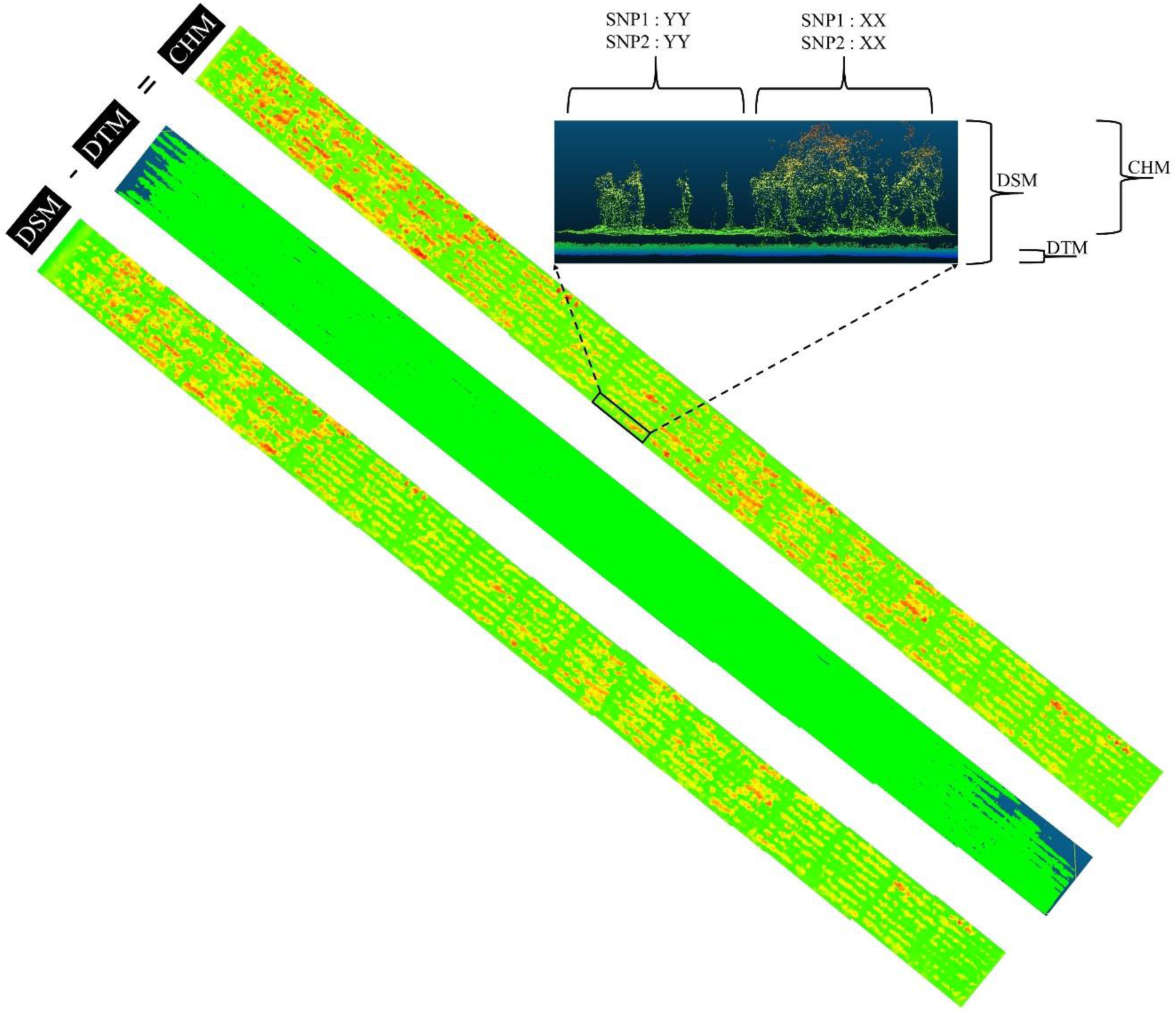
Illustrations and comparisons of canopy height measurements for the HIF-plots with favorable alleles (XX:XX; SNP1:SNP2) versus unfavorable alleles (YY:YY; SNP1:SNP2) using point cloud data.

**Fig. S2.**
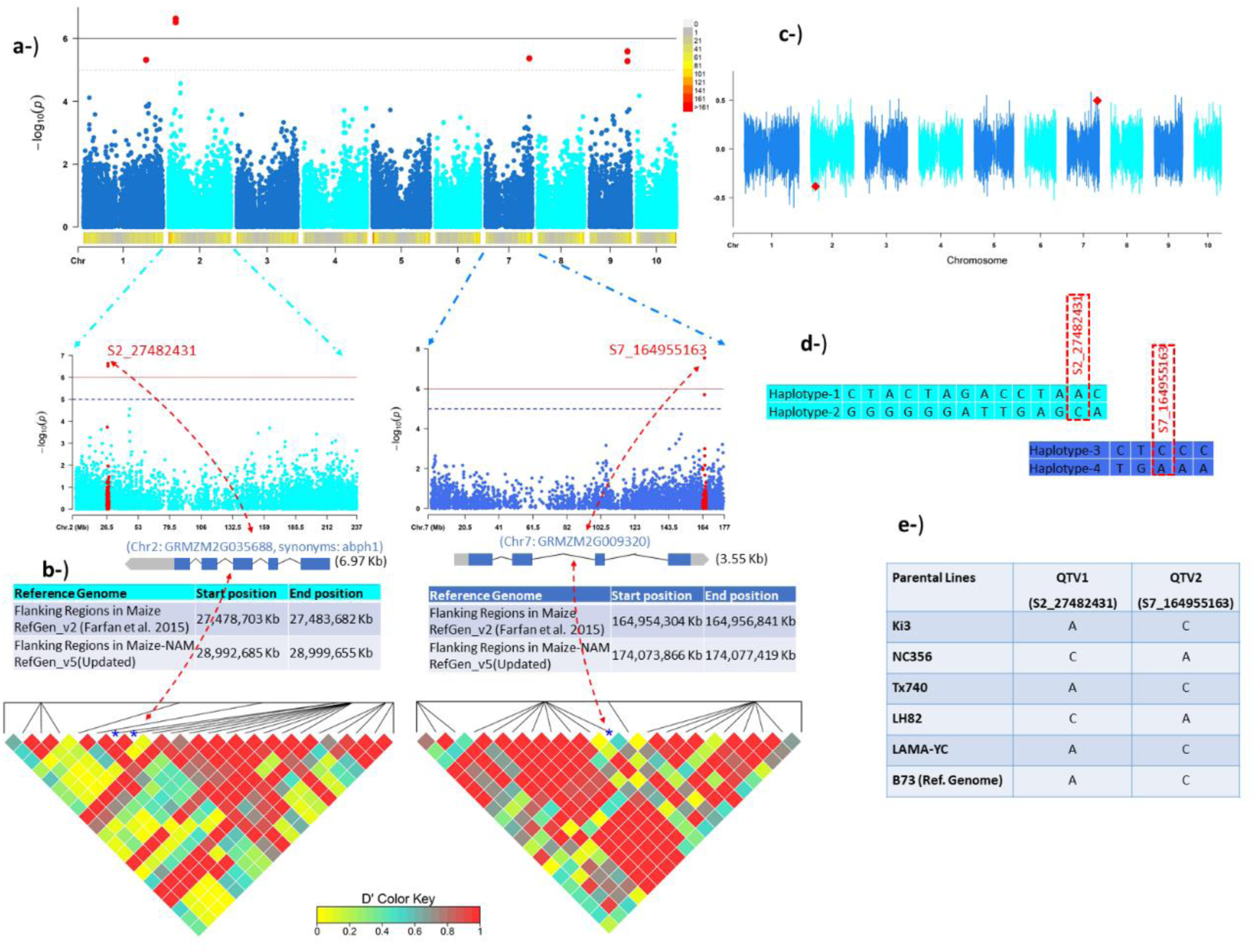
Manhattan plot showing SNP1 and SNP2 used in this study. (a) The two SNPs were discovered in Manhattan plot when plant height was included as a covariate while yield was used as response in the statistical model; SNPs positions updated from maize reference genome version 5 were used to find genes and flanking regions. (b) Linkage disequilibrium (LD) blocks of two SNPs with D prime values. (c) SNP effects of two SNPs (d) Polymorphic SNPs colocalized in LD blocks and haplotype variants based on two SNPs and (e) segregations of two SNPs in parental genotypes, advanced populations used in this study as follows: [LAMA (recurrent parent) x LH82], [Ki3 x NC356 (recurrent parent)], [Ki3 (recurrent parent) x NC356] and [Tx740 (recurrent parents) x NC356]. e or paste legend here. Paste figure above the legend.

**Fig. S3.**
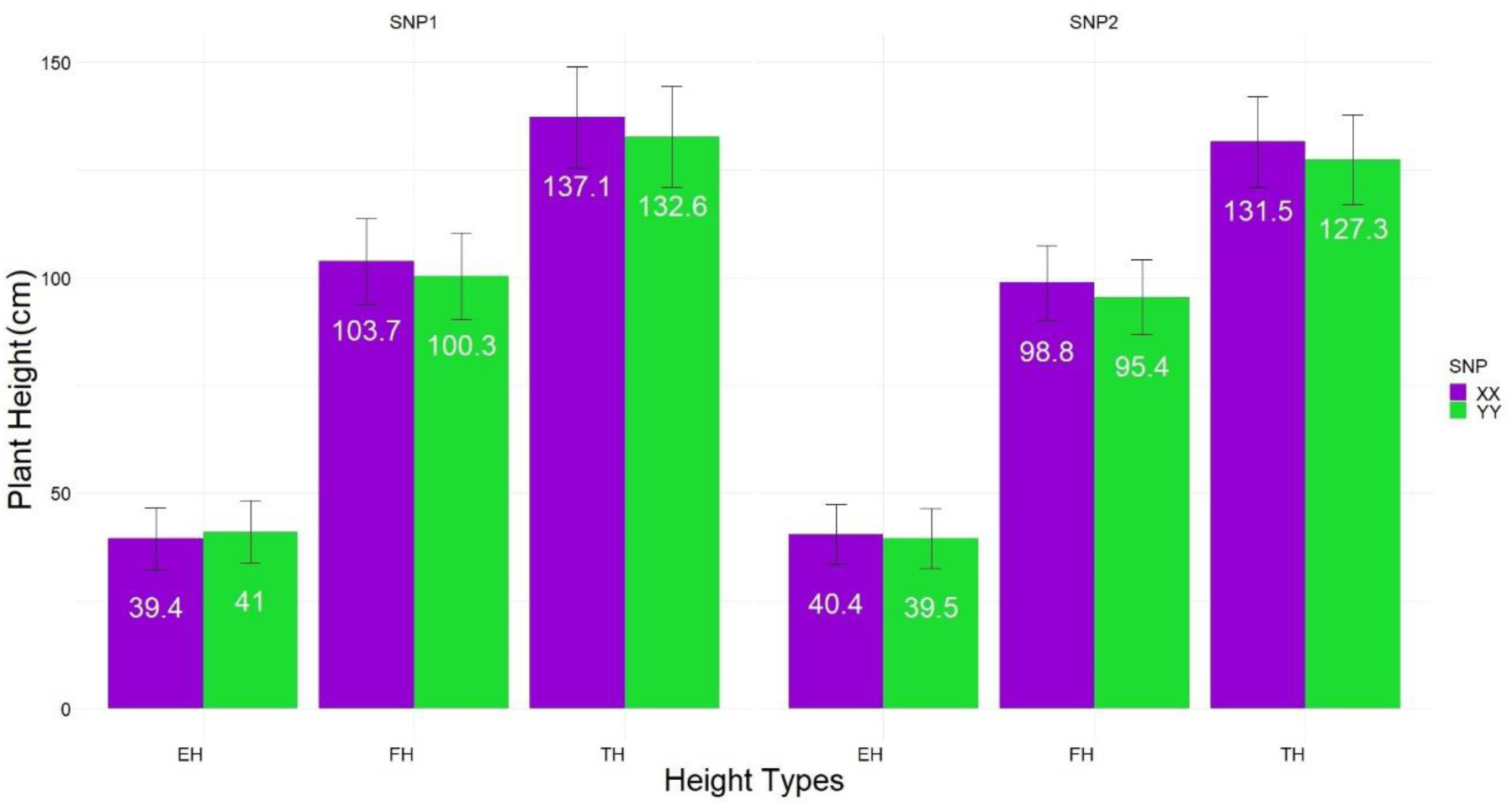
The BLUEs of SNP1 (left) and SNP2 (right) obtained by Equation [2] demonstrating that favorable alleles (XX) contributed consistent taller height for TH and FH, but not for EH. † SNP1 was fixed as XX while SNP2 was segregating to be tested, SNP2 was fixed as XX while SNP1 was segregating to be tested in Equation [2]

**Fig. S4.**
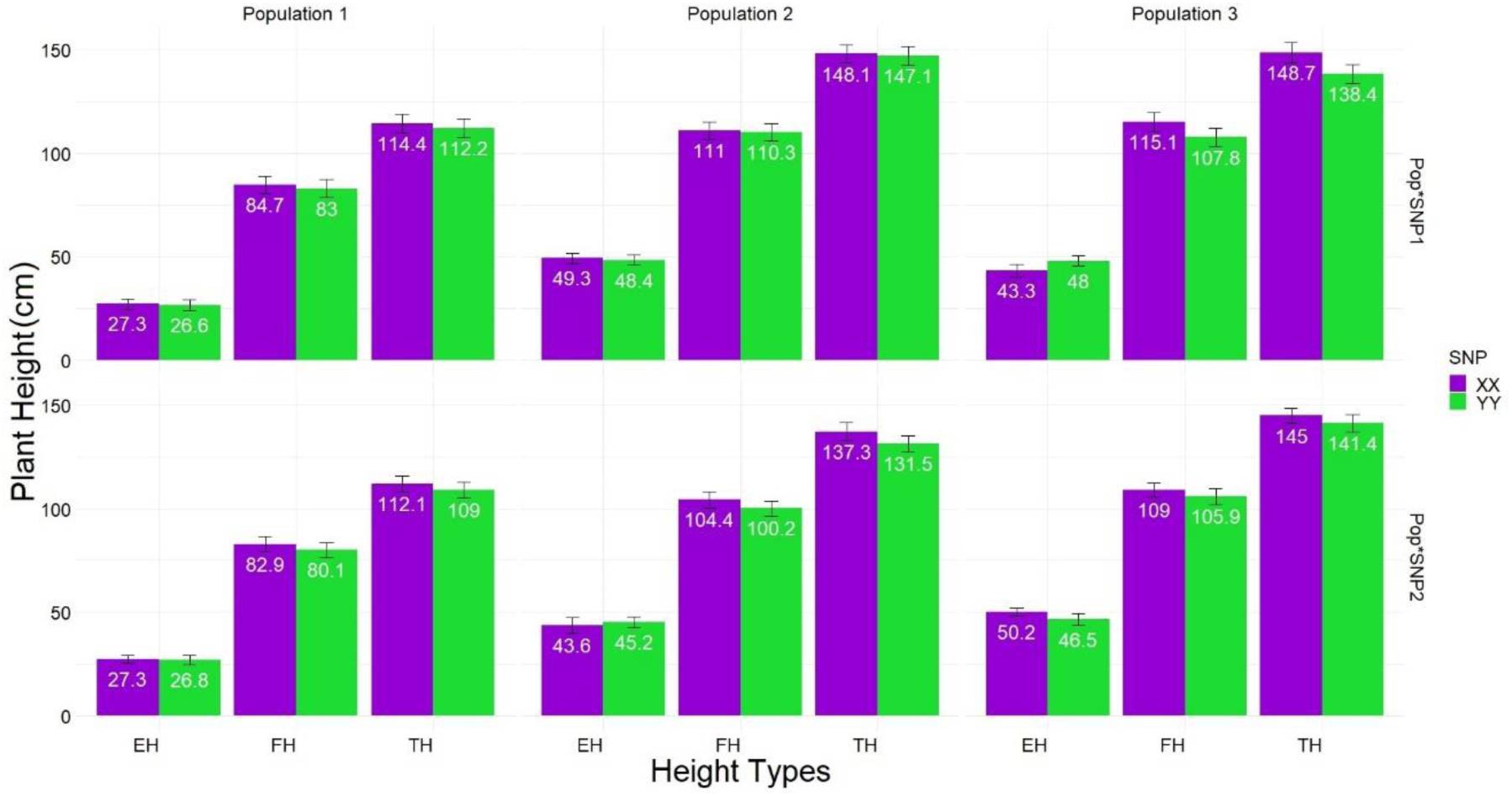
Interaction of SNP1*populations (top) and SNP2*populations (bottom) obtained by Equation [2] shows the largest differences for tassel height. † SNP1 was fixed as XX while SNP2 was segregating to be tested, SNP2 was fixed as XX while SNP1 was segregating to be tested in Equation [2]

**Fig. S5.**
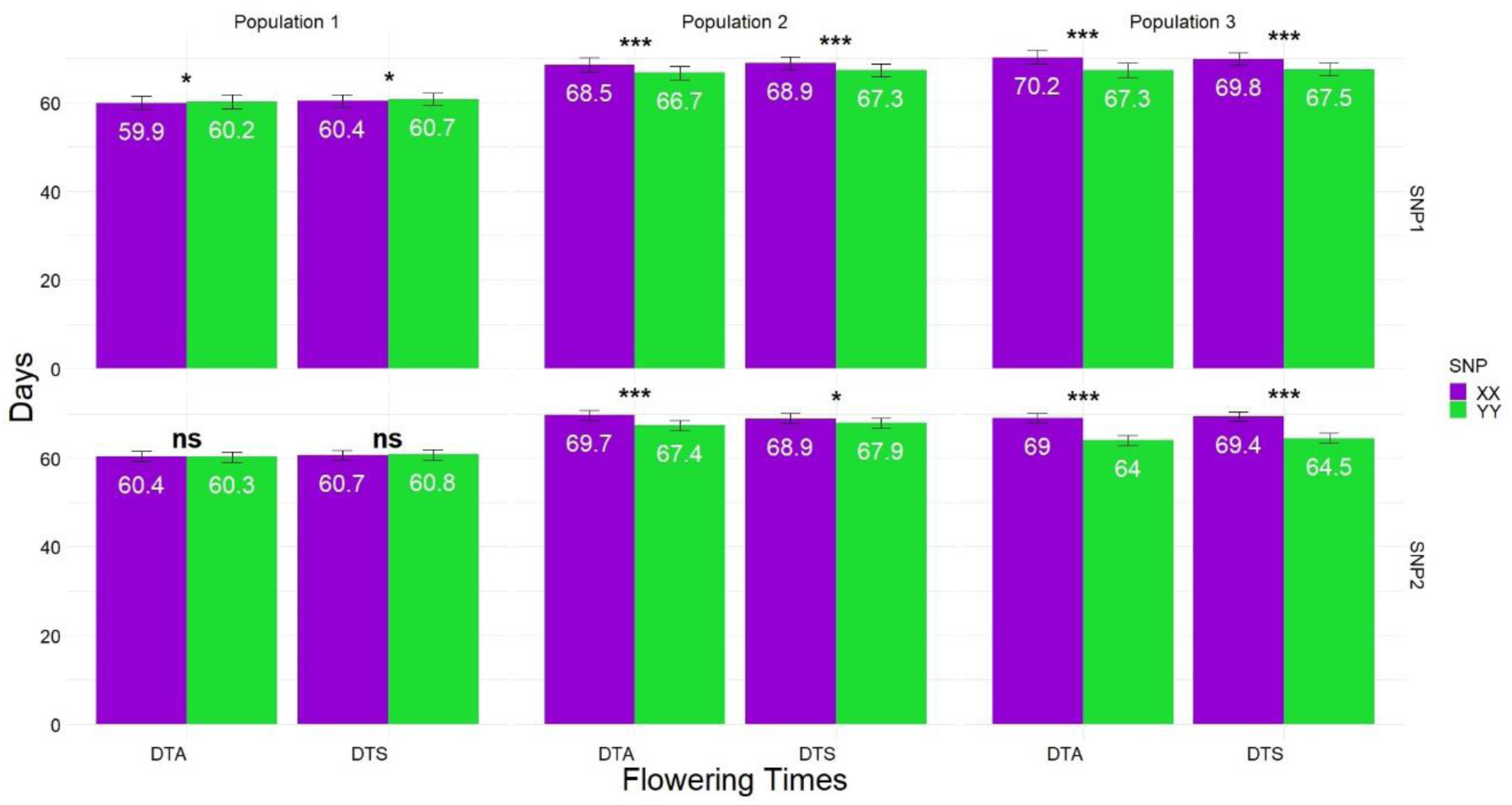
SNP1- and SNP2-population interactions for flowering DTA and DTS in Equation. [2] demonstrating a much larger effect size on population 3. Each call of SNP1 and SNP2 were orthogonally contrasted within population for DTA and DTS. † *, **, *** are the significance level of 0.05, 0.01 and 0.001 respectively.

**Fig. S6.**
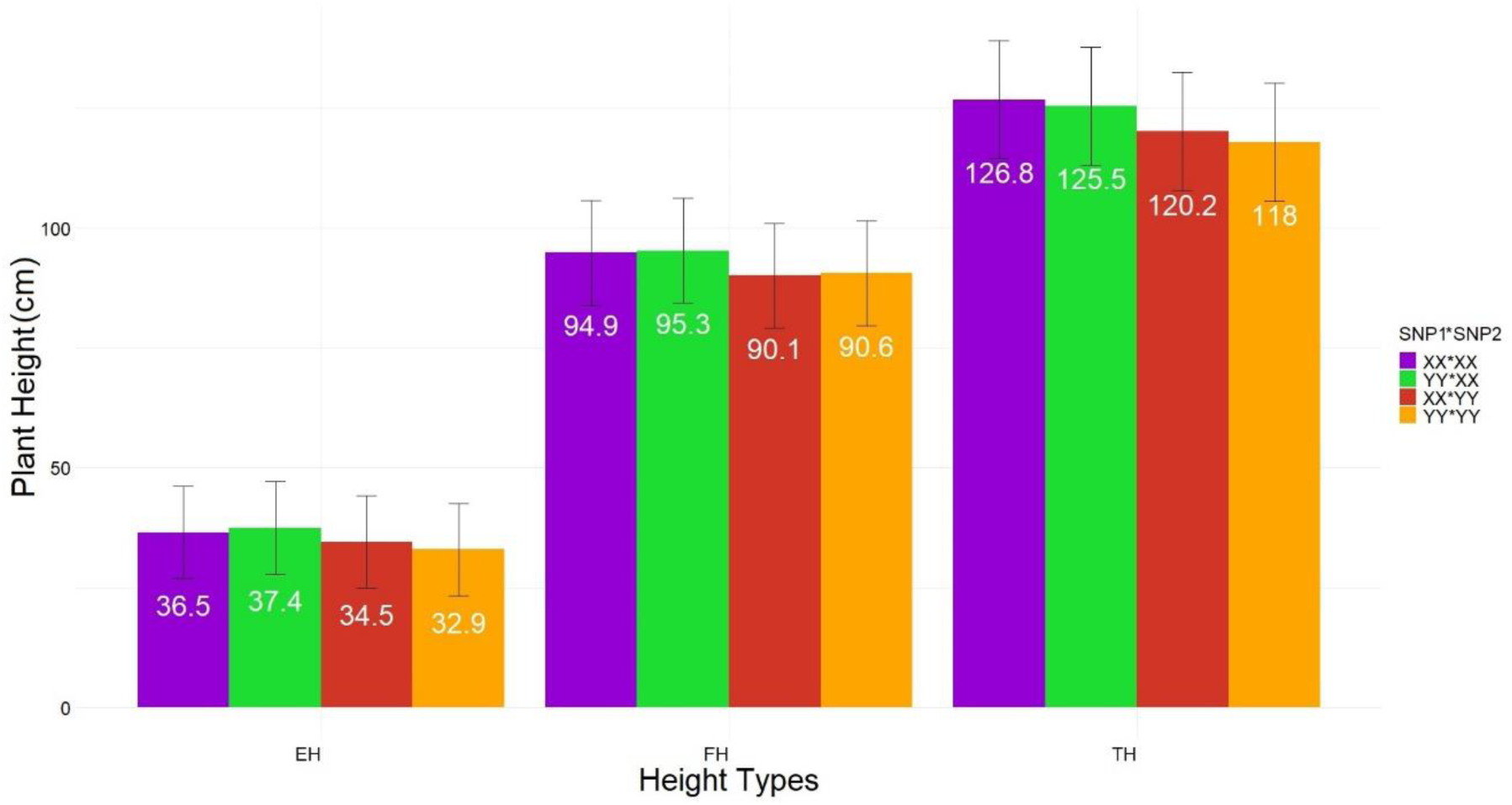
Interaction of SNP1-SNP2 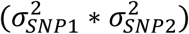 estimated by Equation [3] for each ruler height measurements.

**Fig. S7.**
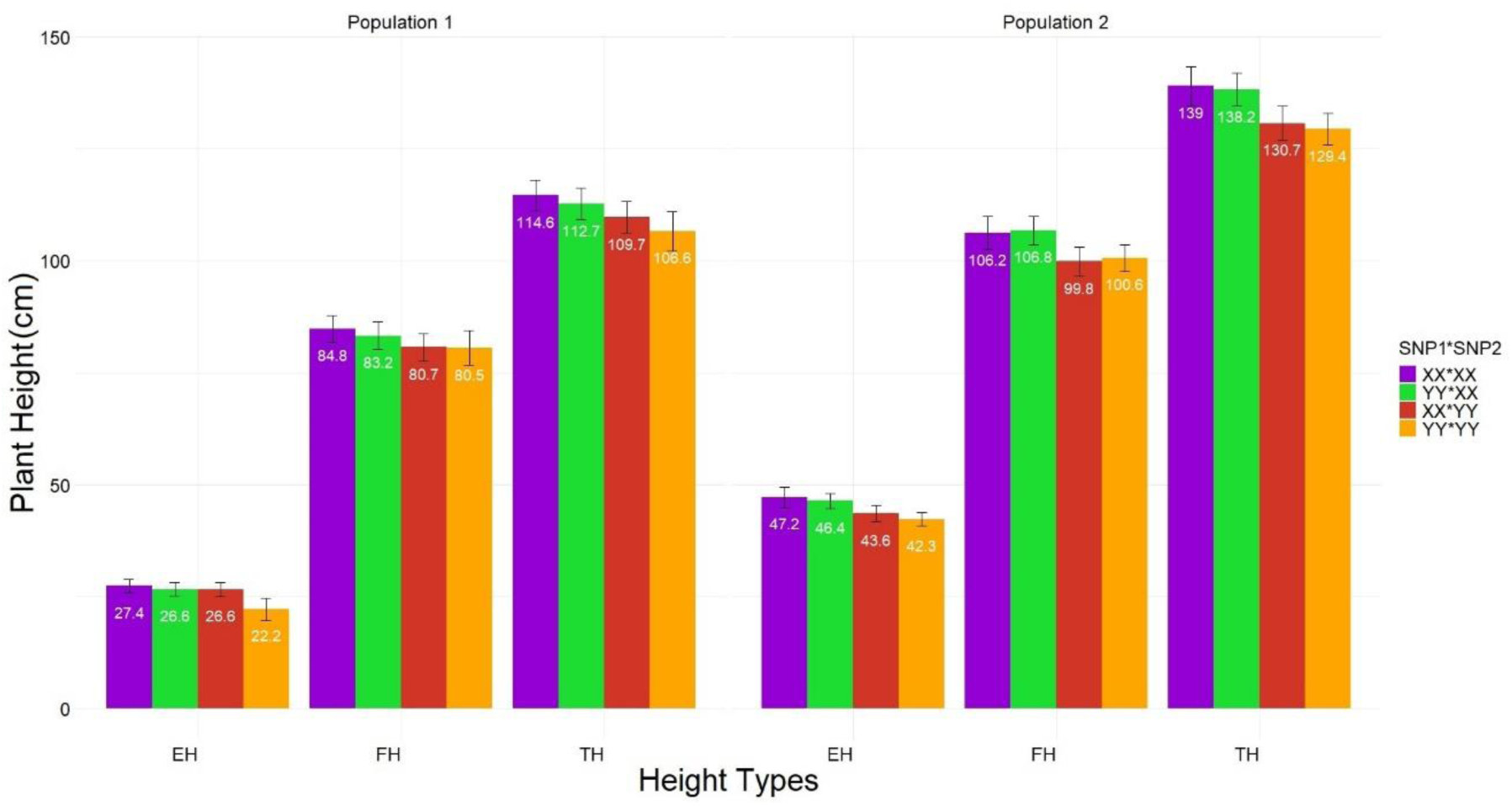
Combined interaction of SNPs with populations 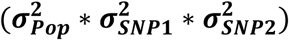.

**Fig. S8.**
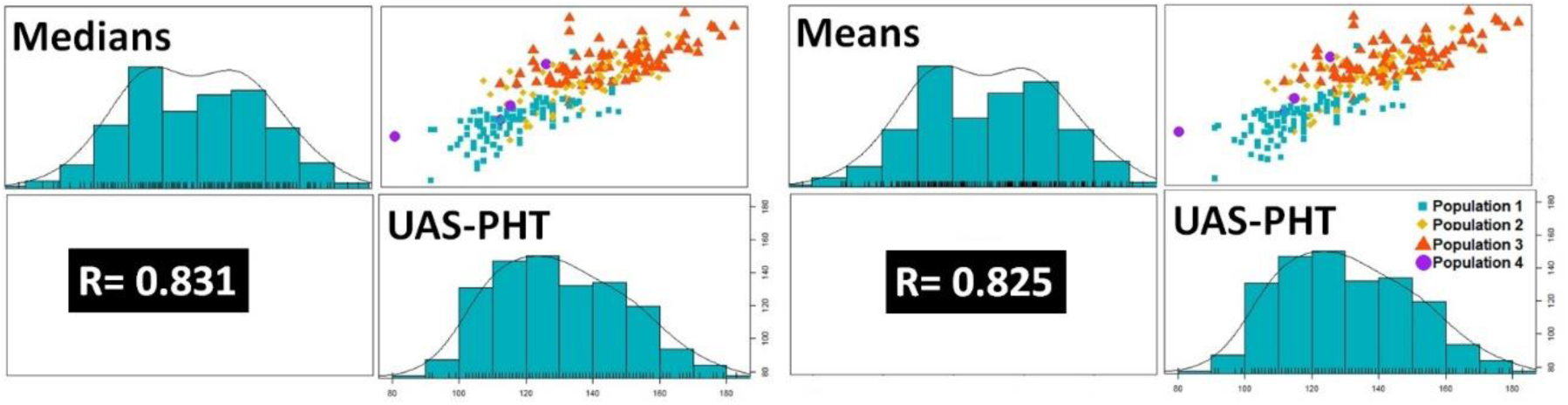
the Pearson correlations (R) between UAS-PHT with means (right) and medians (left) of HIF plots.

**Table S1.** Sources of variance explained by each compoenets in Equation [2] and repeatability for SNP1 (tob) and SNP2 (bottom).

| Variance component<br>(Random effect) | Percentage of variation explained by each variable component for each type of height measurement by ruler |  |  |
| --- | --- | --- | --- |
|  | <u>TH</u> | <u>FH</u> | <u>EH</u> |
| Population | 71.2 | 65.8 | 80.9*** |
| SNP1 | 1.0 | 0.6 | 0.1*** |
| Population*SNP1 | 1.8 | 0.8 | 0.1*** |
| Row | 1.4 | 2.1 | 0.7*** |
| Range | 5.4*** | 7.0*** | 2.9*** |
| Replication | 7.2 | 8.0 | 2.8 |
| Residual | 11.9 | 15.7 | 12.4 |
| **Total variation in number** | 482.4 | 363.4 | 343.1 |
| **Repeatability (R)** | 0.92 | 0.89 | 0.93 |
|  | <u>TH</u> | <u>FH</u> | <u>EH</u> |
| Population | 65.1*** | 77.5 | 69.8 |
| SNP2 | 0.4*** | 0.3*** | 0.1*** |
| Population*SNP2 | 0.1*** | 0.1*** | 0.8 |
| Row | 0.3*** | 0.4*** | 0.3 |
| Range | 9.7*** | 7.1*** | 5.7** |
| Replication | 11.9 | 6.7 | 7.0 |
| Residual | 12.5 | 7.8 | 16.1 |
| **Total variation in number** | 594.2 | 720.5 | 244.4 |
| **Repeatability (R)** | 0.91 | 0.95 | 0.90 |

**Table S2.** Variance component percentage estimates of ruler height from Equation [3].

| Variance component<br>(Random effect) | Percentage of variation explained by each variable component for each type of height measurement by ruler |  |  |
| --- | --- | --- | --- |
|  | **TH** | **FH** | **EH** |
| Population | 63.8*** | 75.4*** | 78.2*** |
| SNP1 | 0.5*** | 0.1*** | 0.1*** |
| Population*SNP1 | 0.2*** | 0.4*** | 0.8*** |
| SNP2 | 5.0*** | 2.3 | 0.3*** |
| Population*SNP2 | 0.8*** | 0.2*** | 0.4*** |
| SNP1*SNP2 | 0.1*** | 0.1*** | 0.6*** |
| Population*SNP1*SNP2 | 0.1*** | 0.1*** | 0.1*** |
| Replication | 3.4 | 2.2*** | 1.0 |
| Row | 1.4*** | 1.1*** | 0.2*** |
| Range | 9.9** | 7.0*** | 3.5*** |
| Residual | 14.9 | 11.1 | 14.8 |
| **Total variation in number** | 467.8 | 515.8 | 299.0 |
| **Repeatability (R)** | 0.90 | 0.93 | 0.91 |

